# PerturbAtlas: A Comprehensive Atlas of Public Genetic Perturbation Bulk RNA-seq Datasets

**DOI:** 10.1101/2024.07.28.605482

**Authors:** Yiming Zhang, Ting Zhang, Gaoxia Yang, Zhenzhong Pan, Min Tang, Yue Wen, Ping He, Yuan Wang, Ran Zhou

## Abstract

Manipulating gene expression is crucial for understanding gene function, with high-throughput sequencing techniques such as RNA-seq elucidating the downstream mechanisms involved. However, the lack of a standardized metadata format for small-scale perturbation expression datasets in public repositories hinders their reuse. To address this issue, we developed PerturbAtlas, an add-value resource that re-analyzes publicly archived RNA-seq libraries to provide quantitative data on gene expression, transcript profiles, and alternative splicing events following genetic perturbation. PerturbAtlas assists users in identifying trends at the gene and isoform levels in perturbation assays by re-analyzing a curated set of 122,801 RNA-seq libraries across 13 species. This resource is freely available at https://perturbatlas.kratoss.site as both raw data tables and an interactive browser, allowing searches by species, tissue, or genomic features. The results provide detailed information on alterations following perturbations, accessible through both forward and reverse approaches, thereby enabling the exploration of perturbation consequences and the identification of potential causal perturbations.

**GRAPHICAL ABSTRACT:** 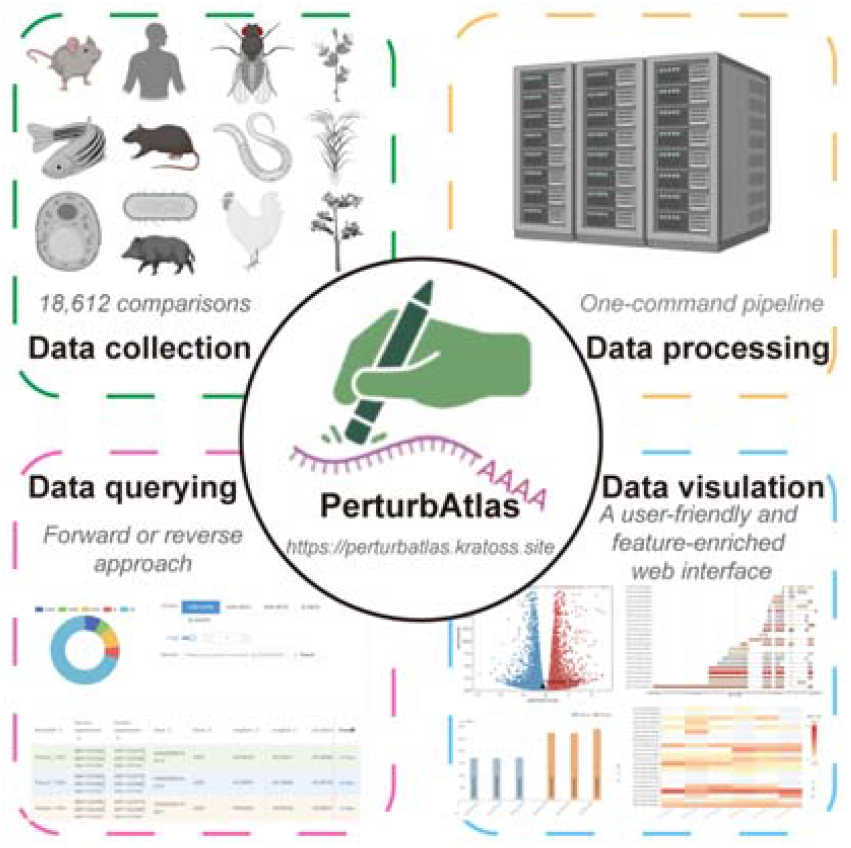

## INTRODUCTION

Gene expression manipulation is essential for understanding gene function, modifying cellular behaviors, and advancing therapeutics(1,2). RNA sequencing (RNA-seq) is widely used to explore the effects of gene perturbations and establish direct links between perturbed genes and their downstream consequences at the gene, transcript and exon levels(3-5). However, despite the increasing availability of gene perturbation RNA-seq data, reusing these public datasets is challenging. One major obstacle is the lack of a standardized metadata format for small-scale perturbation expression datasets in repositories like Sequence Read Archive (SRA)(6) or European Nucleotide Archive (ENA)(7). Additionally, the preprocessing steps, such as data alignment and quantification, are difficult for researchers without high-performance computing resources. Although repositories like GEO(8) or ArrayExpress(9) provide processed gene matrices, differences in gene annotation and normalization methods across studies impede the effective utilization of these datasets. Furthermore, GEO and ArrayExpress often lack in-depth information, such as transcript-level expression and alternative splicing events, further hindering the reuse of these datasets.

So far, several web-based tools are available that provide integrated interfaces for exploring genetic perturbation data. These tools include GPA(10), KnockTF(11), PertOrg1.0(12), and GPSAdb(13). However, GPA primarily focuses on collecting data from microarrays and is currently inactive. GPSAdb and KnockTF databases focus specifically on microarray datasets or RNA-seq data related to gene perturbation assays, limited to species such as human, maize, Arabidopsis or mice. KnockTF exclusively targets transcription factor (TF) genes and does not cover non-TF genes. PertOrg1.0 has been developed to collect perturbation datasets from microarrays and RNA-seq in the GEO. However, it lacks the inclusion of human samples as well as datasets that are not available in GEO but in other repositories such as SRA or ENA. Additionally, the use of the shiny framework in constructing GPSAdb may result in decreased efficiency under high levels of concurrency, and the static visualization of results may not be intuitive and user-friendly. PertOrg1.0 presents results based on a uniform criterion without providing greater freedom and flexibility for users, and it restricts the presentation to a maximum of 1000 genes when the total number of genes exceeds that threshold. Furthermore, these existing databases focus solely on gene expression alterations post-perturbation, neglecting other in-depth information such as different expression of transcripts and alternative splicing regulation. Overall, these tools have limitations in terms of their study coverage, representation of species and diversity of quantitative results (refer to table 1), as well as their efficiency and user-friendliness in the web interface.

In response to this, we meticulously collected and manually annotated all 10,700 perturbation studies from NCBI, EBI, and ENCODE up until February 2023, retaining 7,778 studies with at least one biological replicate for each perturbation assay. This comprehensive dataset comprises 122,801 samples and includes knock-out, knock-down, knock-in, mutation, and over-expression experiments based on RNA-seq data. A total of 257.51T FASTQ files were downloaded and processed with updated gene annotations to re-quantify gene- and transcript-level expression, as well as alternative splicing events. The result is an intuitive and interactive database called PerturbAtlas, which significantly surpasses existing databases in terms of data quantity and species diversity.

## MATERIAL AND METHODS

### Data collection and curation

We systematically collected raw metadata from perturbation datasets using search keywords such as ‘knockdown/-down/ down’, ‘knockout/-out/ out’, ‘overexpression/-expression/ expression’, ‘si/shRNA’, ‘knockin’, ‘dCas9’ and ‘CRISPR-Cas9/ Cas9’ on NCBI/GEO and EBI/ArrayExpress. This process was facilitated by an automated crawler developed in the Go programming language, with the source code available at https://github.com/ygidtu/sra. A total of 116,751 terms were initially collected, resulting in 10,700 studies for further analysis. We manually curated the metadata of each study, including the subject being studied and the type of perturbation targets, retaining only datasets with biological replicates per comparison condition. Datasets lacking information on gene perturbation or tissue/cell line, as well as those derived from underrepresented species with fewer than 10 studies, were discarded, resulting in the retention of datasets from 13 species with varying perturbation assays (Figure 1A). Subsequently, a total of 7,778 studies, comprising 122,801 samples and 18,612 pairs of comparisons derived from sorted cells, cell lines, or tissues (Figure 1B), were used to generate PerturbAtlas.

**Figure 1.**
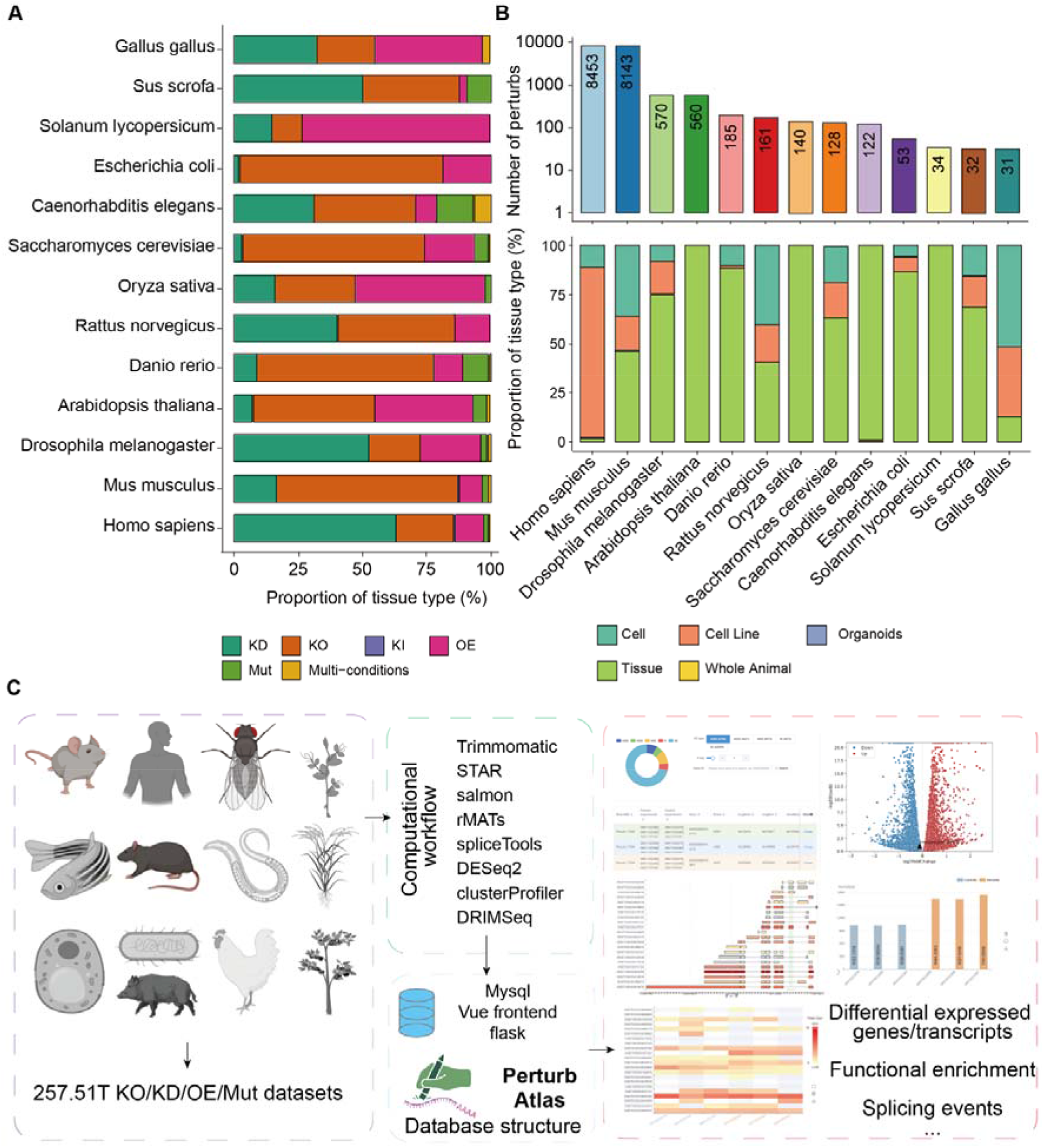
Overview of PerturbAtlas and its Data Composition. **(A)** Distribution of perturbation categories across 13 species in PerturbAtlas. The colors represent various genetic perturbation types: Knockdown (KD), Knockout (KO), Knockin (KI), Overexpression (OE), Mutation (Mut), and Multi-conditions. **(B)** Number of perturbation assays and tissue type distribution. The top panel shows the total number of perturbation assays for each species, with numbers indicating the total assays per species. The bottom panel displays the proportion of different tissue types used in the assays for each species. **(C)** Computational workflow and database structure of PerturbAtlas. The diagram illustrates the pipeline for processing raw data, detailing the steps from raw data collection (257.51T raw data) to the generation of PerturbAtlas. The database structure integrates various analyses including differential expression of genes/transcripts, functional enrichment, and splicing events, using MySQL, Vue frontend, and Flask backend.

### Data preprocess

#### Alignment and quantification

All RNA-seq libraries were analyzed for gene and transcriptomic expression, as well as alternative splicing events, as illustrated in Figure 1C. The complete pipeline, including all analysis scripts, is available at https://github.com/ygidtu/perturb_processing. Briefly, low-quality reads were trimmed using Trimmomatic v0.39(14) with the parameters ILLUMINACLIP:adapters.fa:2:30:10, LEADING:3, TRAILING:3, SLIDINGWINDOW:4:15, and MINLEN:25. The trimmed reads were then aligned to the corresponding genome using STAR v2.7.10b(15) with default parameters to generate BAM files as well as the table named ReadsPerGene.out.tab. Gene counts and transcripts per million (TPM) were obtained using Salmon v1.10.1(16) with –numBootstraps 100. Functional enrichment analyses, such as Gene Ontology (GO)(17) and Kyoto Encyclopedia of Genes and Genomes (KEGG)(18), were performed by the clusterProfiler package(19), utilizing the OrgDb from Bioconductor. If the OrgDb was not available, representative annotations were downloaded and processed with the GO.db package to generate GO terms for enrichment analysis.

#### Different gene and transcript expression analysis

Differentially expressed genes were identified using DESeq2(20), which performed normalization and differential expression testing on the raw read counts for each gene. The final tables were exported and used to prepare the database. For differential transcript usage (DTU) analysis, the transcription expression data and bootstrap results generated by Salmon were processed using DRIMSeq(21) with default parameters. The resulting table, which includes a p-value for each transcript and the transcript proportions of a given gene, was then incorporated into the database.

#### Alternative splice analysis

When experimental information was not available, the strandedness of the RNA-seq libraries was inferred from the ReadsPerGene table generated by STAR. Specifically, if the ratio of reads aligned to the stranded or reverse-stranded orientations exceeded 80%, the library was classified as fr-secondstrand or fr-firststrand, respectively. Libraries not meeting this criterion were classified as unstranded. Subsequently, alternative splicing (AS) events were identified and quantified using rMATS(22) in paired mode, accounting for strandedness, utilizing BAM files generated by STAR. Furthermore, the summary statistics and potential functions of AS results were analyzed using spliceTools. Collectively, the final table was incorporated into the database.

### Database implementation

The current iteration of PerturbAtlas was developed using Postgres 15.3 (https://www.postgresql.org/) and is hosted on a Linux-based Caddy server (https://caddyserver.com/). Server-side scripting was implemented using Python Flask 3.0.2 (https://flask.palletsprojects.com/en/). For the interactive interface, Vue 3.4.29 (https://vuejs.org/) and ElementPlus v2.3.1 (https://element-plus.org/) were utilized to ensure a user-friendly experience. To provide graphical visualizations, ECharts 5.5.0 (https://www.echartsjs.com/) and matplotlib 3.9.0 (https://matplotlib.org/) were integrated as the visualization frameworks. Analysis-related functions rely on R version 4.2.2, and the entire environment is set up using Docker. For optimal display and functionality, a modern web browser that supports the HTML5 standard, such as Firefox, Google Chrome, Safari, Opera, or IE 9.0+, is recommended. The PerturbAtlas database is freely accessible to the research community through the provided web link, and no registration or login is required to access its features.

## RESULTS

### Overview

The user interface of PerturbAtlas is depicted in Figure 2. PerturbAtlas enables dataset exploration through both forward and reverse approaches. In the forward approach, users can search for data of interest using a public accession number, the target of perturbation, the name of the cell, tissue, or organ, or a combination of these parameters (Figure 2A). The eligible results will be displayed, including information on treatment conditions and disease status, if available. Upon selecting a specific term, users will receive a detailed description curated from the original project description, assisting them in assessing the suitability of the data for their specific needs. Once a specific dataset is selected, PerturbAtlas offers four modules for further data exploration: “Differentially Expressed Genes”, “Enrichment Analysis Results”, “Alternative Splicing”, and “Differentially Expressed Transcripts” (Figure 2C and D). Each module provides detailed insights into the dataset, facilitating comprehensive analysis. Moreover, PerturbAtlas integrates various results to offer a holistic view. For instance, if a gene of interest is identified as significantly differentially expressed in the current assay, the database also provides information on alternative splicing events and transcript expression levels. This integration helps users plan subsequent steps in their research. To maximize utility and allow users to set their own thresholds, all data in PerturbAtlas is presented in an unfiltered form. Users can apply flexible criteria to filter the results according to their specific needs.

**Figure 2.**
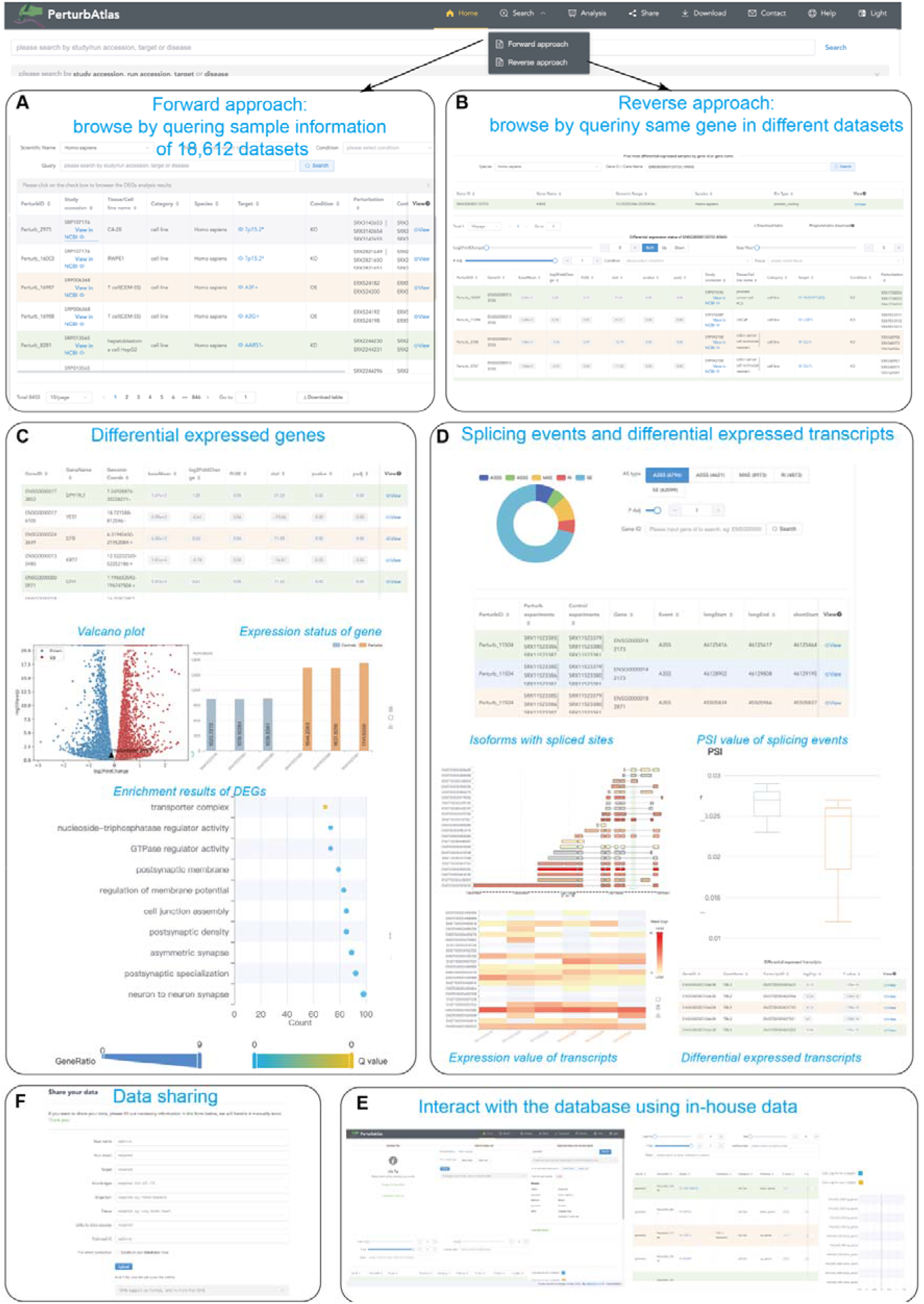
User Interface and Functionality of PerturbAtlas. (A) In the forward approach, users can browse through 18,612 datasets by querying sample information using public accession numbers, perturbation targets, or the names of cells, tissues, or organs. (B) In the reverse approach, users can explore the same gene across different datasets, adjusting parameters such as fold change and adjusted p-value to refine results and identify potential up-regulators. (C) The Differentially Expressed Genes module provides insights into differentially expressed genes, including volcano plots, gene expression statuses, and enrichment results for DEGs. (D) The Splicing Events and Differentially Expressed Transcripts module offers detailed information on splicing events, isoforms with spliced sites, PSI values of splicing events, and the expression values of transcripts. (E) The data sharing module allows users to upload their own datasets for sharing and future integration into PerturbAtlas. (F) PerturbAtlas offers an enrichment-based approach that allows users to interact with the database, further identifying potential regulators of the gene set.

**Figure 3.**
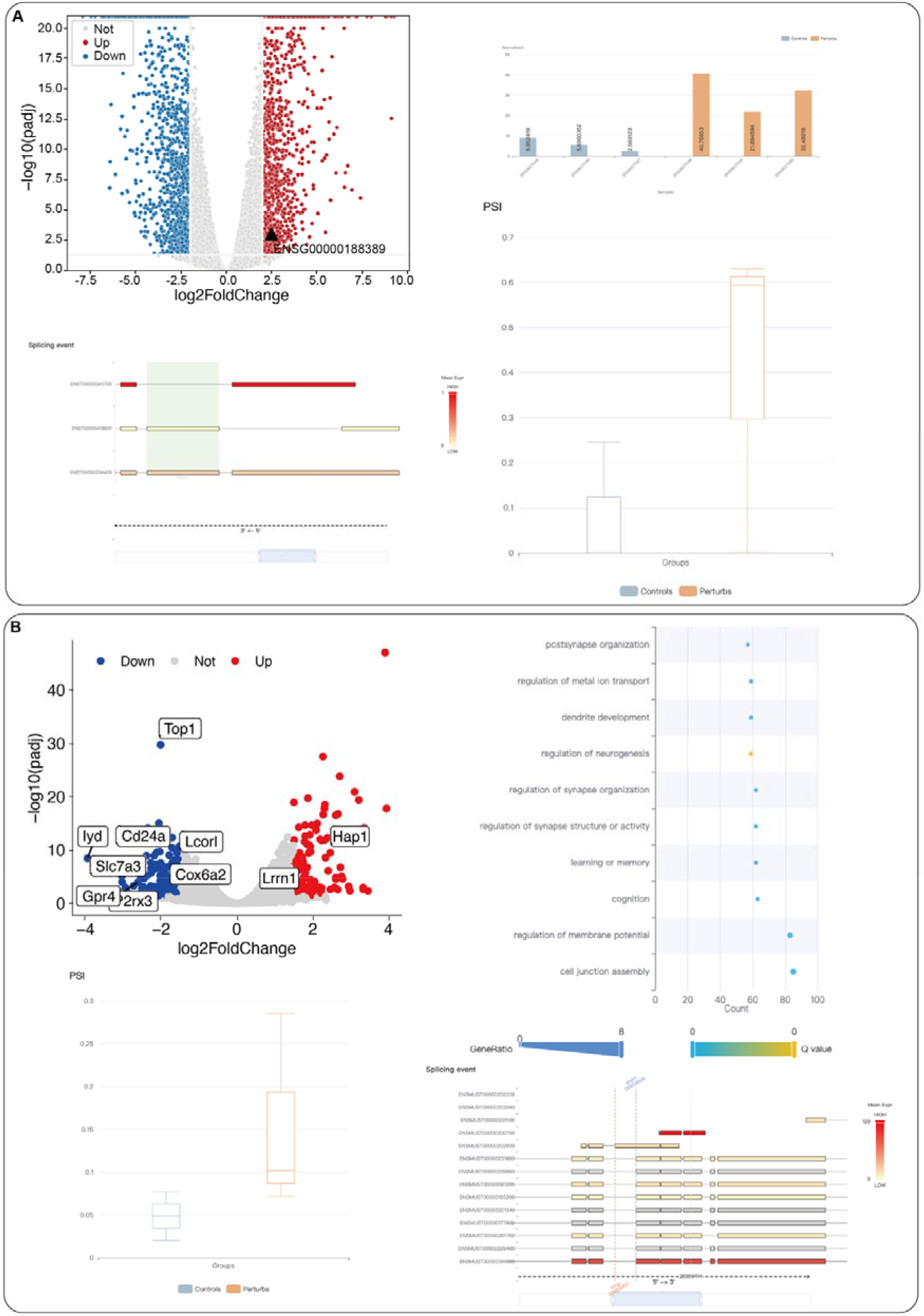
The example of PerturbAtlas usage. (A) The volcano plot illustrates the differentially expressed genes in mouse cortical neural tissue before and after *Top1* knockout. Red dots indicate significantly upregulated genes, blue dots indicate significantly downregulated genes, and grey dots represent genes with no significant changes. The dot plot highlights the enrichment of GO pathways for differentially expressed genes following *Top1* knockout. The box plot shows the PSI values of the Syngap1 gene Exon11 A3SS event before and after Top1 knockout. Additionally, the transcript structure plot reveals the A3SS event site of the *Syngap1* gene E11, with transcript colors indicating average expression levels. (B) The volcano plot displays differentially expressed genes in the human non-small cell lung cancer H358 cell line following *SMARCA4* knockout. Red dots represent significantly upregulated genes, blue dots represent significantly downregulated genes, and grey dots indicate genes with no significant changes. The bar chart shows the expression levels of *PDCD1* before and after *SMARCA4* knockout. The transcript structure plot illustrates the ES event site of the *PDCD1* gene E3, with colors representing average expression levels. The box plot displays the PSI values of the *PDCD1* gene Exon3 ES event before and after *SMARCA4* knockout.

In the reverse approach, users can investigate the differential expression of their gene of interest across all datasets within a specific species (Figure 2B). PerturbAtlas provides metadata of the relevant perturbation terms, along with differential expression details such as expression levels, fold changes, and adjusted p-values for the target gene. Users can refine their search by adjusting various parameters, including fold change and adjusted p-value, to narrow down the results and identify potential up-regulators of the target gene. Once a perturbation term of interest is selected, an interactive interface will be provided, encompassing gene and transcript expression, as well as alternative splicing regulation if available.

### Download and Share

PerturbAtlas offers unrestricted access to all analysis results, which users can easily download from the Download page (Figure 2F). This page provides extensive information on perturbation datasets, including dataset details, differential expression analysis results, gene and transcript expression profiles, and alternative splicing results. Despite efforts to collect bulk RNA-seq data related to perturbations up until February 2023, some datasets lacking explicit perturbation keywords or sample descriptions may have been missed. To ensure comprehensiveness, users are encouraged to submit their own datasets for future integration. Through the “Submit” page, users can provide details such as perturbed genes, perturbation types, tissue types, data types, organisms, and links to their data sources. Upon submission, the data will be evaluated and processed according to standard procedures, and the dataset will be included in future releases of PerturbAtlas, enhancing its content and utility.

### Database functionality

PerturbAtlas offers an interactive analysis module that allows users to integrate their own datasets, whether they involve genetic manipulation (Figure 2E). Users can upload a file containing gene names and corresponding fold change values. PerturbAtlas will then perform a Gene Set Enrichment Analysis (GSEA) using signatures from each perturbation term. This analysis enables users to identify genes that, after perturbation, exhibit similar transcriptional responses to those in the uploaded data. This functionality provides deeper insights into the effects of various assays, such as drug treatments, and their transcriptional impacts. Additionally, PerturbAtlas allows users to upload a list of genes of interest to determine which perturbation terms most significantly enrich these genes, further identifying potential regulators of the gene set. In summary, PerturbAtlas offers an enrichment-based approach that allows users to interact with the database using their in-house data. PerturbAtlas performs the analysis without storing the uploaded data or collecting user information, ensuring user privacy and data security.

### Exemplified use-cases

#### Case study 1

In this case, we aimed to explore the potential regulation of PDCD1, a gene associated with immune therapy by using the reverse approach page of PerturbAtlas. After initiating the search, we applied filters to identify 69 datasets where PDCD1 expression showed significant changes, defined by abs(log2FoldChange) > 2 and p adj < 0.05. We further refined the search to focus on non-small cell lung cancer by using the tissue search box. The results indicated that following *SMARCA4* knockout, PDCD1 expression was significantly upregulated, consistent with reports of promising outcomes in treating *SMARCA4*-deficient tumors with PD-1/PD-L1 inhibitors(23). Additionally, PerturbAtlas revealed exon skipping in the third exon of PDCD1, suggesting potential post-transcriptional regulation affecting PDCD1 function.

#### Case study 2

Before conducting experimental studies on a gene of interest, it is crucial to gather information and generate hypotheses to guide the research. For instance, when studying the *Top1* gene, which encodes a DNA topoisomerase, one could use PerturbAtlas to examine the consequences of *Top1* perturbation. Following the knockout of *Top1*, genes related to synapse organization and neurogenesis, such as *Hap1*, were significantly upregulated, suggesting that *Top1* may regulate neurogenesis through the modulation of these genes in neuronal cells. Alternative splicing is widely found in the brain and is involved in the homeostasis of the neuronal system(24,25), PerturbAtlas also allows users to explore changes in alternative splicing events after perturbation. For example, an A3SS event related to *Syngap1* Exon11 is observed in this sample, potentially leading to nonsense-mediated mRNA decay and consequently affecting gene expression levels (26).

## Conclusion and future directions

PerturbAtlas offers a comprehensive bulk RNA-seq database, enabling users to explore gene expression, transcript profiles, and alternative splicing regulation following genetic perturbation. It facilitates efficient in silico screening across 122,801 RNA-seq datasets, providing researchers with new opportunities to gain deeper insights into the molecular mechanisms and networks governing gene regulation. For biologists, the user-friendly interface supports the testing of biological hypotheses and guides experimental design. For computational users, PerturbAtlas provides ready-to-use expression matrices and perturbation metadata, facilitating tasks such as training large language models for perturbation predictions. Additionally, we have developed a fully automated pipeline for database construction (https://github.com/ygidtu/perturb_processing), which simplifies future maintenance tasks such as adding new datasets and updating reference genomes. Future updates will include new perturbation datasets and data on exon and intron expression levels.

## DATA AVAILABILITY

All scripts used to generate the database inputs are available at https://github.com/ygidtu/perturb_processing. The tool for fetching perturbation terms from NCBI and EBI resources can be accessed at https://github.com/ygidtu/sra. The processed matrix utilized to create the interactive interface is available for download at https://perturbatlas.kratoss.site/#/download.

## SUPPLEMENTARY DATA

Supplementary Data are available at NAR online.

## AUTHOR CONTRIBUTIONS

Ran Zhou: Conceptualization, Funding acquisition, Supervision, Formal analysis, Data curation, Methodology, Validation, Writing—original draft. Yuan Wang: Funding acquisition, Supervision, Writing—review. Yiming Zhang: Conceptualization, Formal analysis, Visualization & editing. Ting Zhang: Data curation. Gaoxia Yang: Data curation. Zhenzhong Pan: Data curation. Min Tang: Data curation. Yue Wen: Data curation. Ping He: Data curation. Yaojia Liu: Data curation.

## ACKNOWLEDGEMENTS

After completing the final manuscript, we utilized ChatGPT (OpenAI, https://chat.openai.com/) to proofread the final draft.

## TABLE AND FIGURES LEGENDS

Table 1 Statistics of the existing perturbation databases. It’s important to note that for a single GEO ID or NCBI Project ID, multiple pair comparisons were counted only once.

